# Prenatal stress programs behavioral pattern separation in adult mice

**DOI:** 10.1101/2022.01.10.475616

**Authors:** S Rajendran, ML Kaci, E Ladeveze, DN Abrous, M Koehl

## Abstract

Stress is an unavoidable condition in human life. Stressful events experienced during development, including *in utero*, have been suggested as one major pathophysiological mechanism for developing vulnerability towards neuropsychiatric and neurodevelopmental disorders in adulthood. One cardinal feature of such disorders is impaired cognitive ability, which may in part rely on abnormal structure and function of the hippocampus. In the hippocampus, the dentate gyrus is a site of continuous neurogenesis, a process that has been recently implicated in spatial pattern separation, a cognitive phenomenon that serves to reduce the degree of overlap in the incoming information to facilitate its storage with minimal interference. We previously reported that adult neurogenesis is altered by prenatal stress allowing us to hypothesize that prenatal stress may possibly lead to impairment in pattern separation. To test this hypothesis, both control (C) and prenatally stressed (PS) adult mice were tested for metric and contextual discrimination abilities. We report for the first time that prenatal stress impairs pattern separation process, a deficit that may underlie their cognitive alterations and that may result in defective behaviors reminiscent of psychiatric illness such as post‐traumatic stress disorder.

## Introduction

Early life adversity experienced as early as during the *in utero* development has repeatedly been associated with a negative impact on offspring’s health and development (Glover, 2011; Van den Bergh et al., 2017; Glover et al., 2018) and identified as a risk factor for a wide range of psychiatric disorders such as depression, anxiety, post‐traumatic stress disorder (PTSD) and stress‐related or impulse control disorders (Lupien et al., 2009; Glover et al., 2018). These abnormalities arise from numerous alterations in brain structures that preferentially target stress/anxiety structures, among which the hippocampus. Indeed, the hippocampus is a key target of the stress response; it expresses high levels of corticosteroid receptors in early development (Meaney et al., 1985) and is thus predicted to be especially vulnerable to the effects of developmental stress. In support of this, meta‐analyses report significant associations between childhood adversity, reduced hippocampal volume and impaired hippocampal function (Calem et al., 2017).

Although informative, studies in humans do not allow extensive investigation into the underlying mechanisms of the impact of prenatal stress, and animal models have been developed. It was thus reported in rodents that prenatal stress, which leads to increased anxiety, anhedonia and severe cognitive dysfunctions (Weinstock, 2008), can produce significant alterations in the adult hippocampus. In particular, we and others have reported in rodent and non‐human primate models that PS adversely affects cell proliferation and neurogenesis in the adult brain (Lemaire et al., 2000; Coe et al., 2003; Mandyam et al., 2008; Zuena et al., 2008; Lucassen et al., 2009; Belnoue et al., 2013; Ortega‐Martinez, 2015). Because adult‐born dentate granule neurons are essential for dentate gyrus information processing and are implicated in emotional regulation and mental disorders (Kheirbek et al., 2012a; Yun et al., 2016), prenatal stress‐induced impairments in adult hippocampal neurogenesis and dentate gyrus function are considered critical pathophysiological factors that negatively affect behavior and leave individuals vulnerable to developing psychiatric illnesses.

Adult neurogenesis is believed to play a crucial role in a number of hippocampal‐dependent cognitive processes, including spatial learning and behavioral pattern separation (Clelland et al., 2009; Creer et al., 2010; Sahay et al., 2011a; Kheirbek et al., 2012a; Nakashiba et al., 2012; Tronel et al., 2012). Pattern separation is a process that is commonly defined as the ability to form separate representations from highly similar, yet slightly different, events or stimuli. It refers to the computational process that, after the integration of all sensory inputs (visual, olfactory, vestibular, auditory somatosensory), separates similar events from each other via an orthogonalization of sensory input information (Rolls & Kesner, 2006; Kesner, 2007). This orthogonalization plays a key role in the match–mismatch process – more orthogonolized representation of the changed environment is likely to facilitate detection of mismatch‐, which should help the brain separate individual events that are part of incoming memories (McAvoy et al., 2015). This process is also believed to facilitate recall, by preserving the uniqueness of a memory representation.

Alterations in spatial and contextual memory abilities (Lemaire et al., 2000; Lee et al., 2011) together with the altered adult neurogenesis observed after prenatal stress thus led us to hypothesize that prenatal stress may alter pattern separation abilities. To address this hypothesis, control and prenatally‐stressed adult male mice were challenged in 2 behavioral tasks allowing to assess behavioral pattern separation: a metric processing task that analyzes the ability to discriminate metric relationships between objects (Goodrich‐Hunsaker et al., 2008; Chen et al., 2012) and a contextual fear discrimination task that tests mice abilities to discriminate two similar contexts (Tronel et al., 2012). We report that prenatally‐stressed mice are impaired in both tasks, indicating that prenatal stress certainly leads to a general alteration of pattern separation independently of the tested modality.

## Materials and Methods

### Animals

Eight weeks old female CD1 mice (Charles River, France) were collectively housed under a 12h light/12h dark cycle (lights on from 8 am to 8 pm) in a temperature‐ (22+/‐3°C) and humidity‐controlled facility. Animals had ad libitum access to food and water. After 10 days of habituation to the housing conditions, females were mated with CD1 males (Charles River, France) following a 3 females/1 male breeding design. Pregnancy was determined by the presence of a vaginal plug checked once a day at the beginning of the light phase. Each female was removed from the breeding couple and individually housed upon plug detection (gestational day GD 0). All procedures and experimental protocols were approved by the Animal Care Committee of Bordeaux (CEEA50) and were conducted in accordance with the European community’s council directive of 24 November 1986 (86/609/ECC) that was in effect at the time of the experiments. All efforts were made to minimize animal suffering, and reduce the number of animals used.

### Gestational stress

Gestational stress was carried out from day 8.5 of gestation until parturition (∼GD18‐GD19). Females were restrained in plastic transparent cylinders (in 50 mL centrifuge tubes 3 cm diameter, 11 cm long from GD8.5 until GD15; in vials 5.3 cm diameter, 10.5 cm long from GD15 on) for 45 min three times per day under bright light during the light cycle (9am, 1pm, 5pm). Control mothers were left undisturbed throughout gestation. After delivery, dams and pups were left undisturbed, except for cage change, until weaning (21st day post‐partum). At weaning, only males originating from litters containing between 9 and 15 pups (mean pups per litter=12) with a nearly equal distribution between males and females were kept and group housed with cagemates originating from different mothers. Two groups were thus constituted: a Prenatally‐Stressed group (PS) and a Control group (C).

### Behavioral assessment

Two different batches of mice were used. The first one was tested in a metric pattern separation test at 4 months of age; the second one was tested in a contextual discrimination task at 5 months of age. In each case, mice were individually housed at least 2 weeks before testing, and in order to avoid “litter effect”, a maximum of 2 mice from the same litter was included into each batch. All the studies were performed during the first half of the light period.

### Metric separation

Behavioral pattern separation was measured in a variation of the metric processing task (Goodrich‐Hunsaker et al., 2008) in which the ability of mice to discriminate the metric relationship between two objects under challenging conditions is tested (Chen et al., 2012). The test took place in an open‐field (L=100 cm; W=50 cm; walls H = 30 cm) containing two distinct objects (10‐15 cm high, 5‐8 cm wide). Objects composing each pair were selected so as to induce similar exploration times in preliminary studies. In the sample trial, mice were placed in the center of the open‐field that contained the two objects placed 20 cm apart, and they were allowed 20 min of free exploration of the objects (Figure 1A). During a 5 min inter‐trial interval they were returned to their homecage and the objects were repositioned at 55 cm (low separation load, easy condition), 45 cm (medium separation load), or 35 cm (high separation load, difficult condition) apart. After the intertrial interval, mice were returned to the open field for a 5 min test trial, during which they were able to reinvestigate the objects in their new arrangement. The time that each mouse spent exploring the two objects was recorded during both the sample and test trials. An investigation ratio was calculated as: (exploration during the 5 min test trial)⁄(exploration during the 5 min test trial + exploration during the last 5 min of the sample trial). Each mouse was tested on each of the object separations, in a counter‐balanced design, during different sessions over the course of three consecutive days.

**Figure 1:**
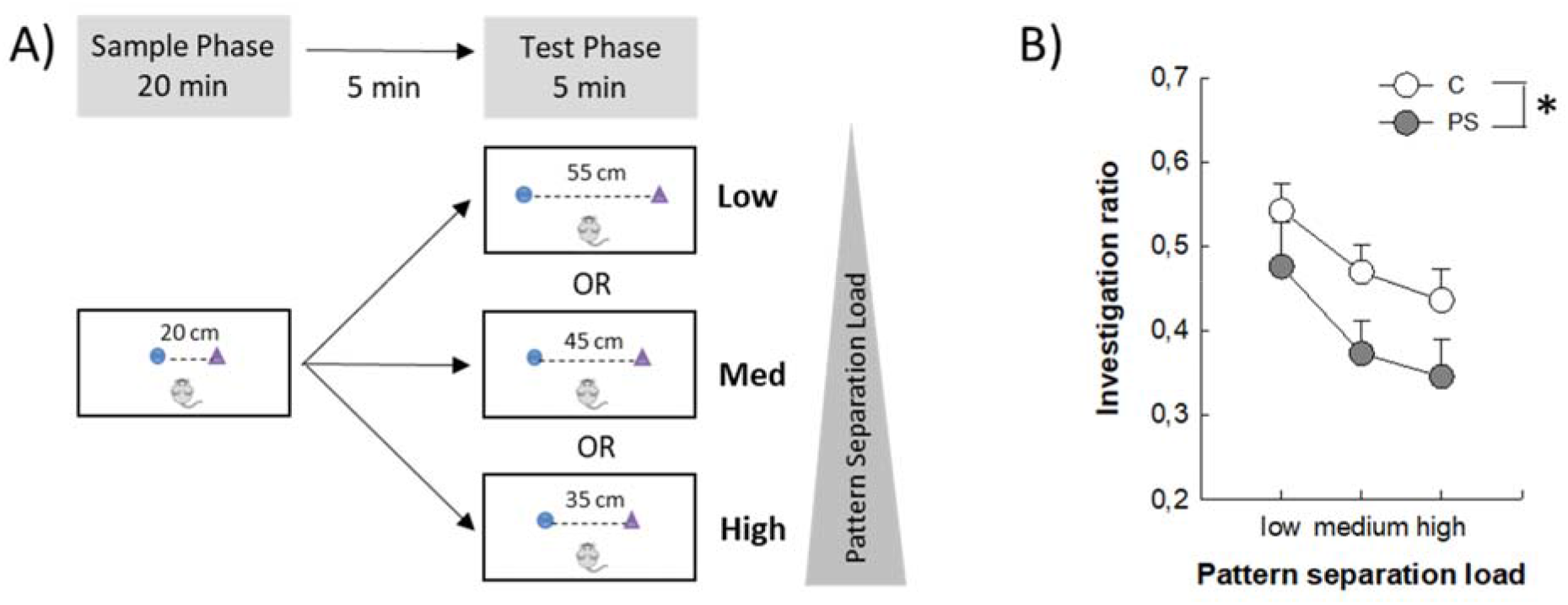
Prenatal stress alters mice abilities to perform metric separation. A) Experimental design of the task. All mice were tested in the three objects arrangements according to a counter‐balanced design over 3 consecutive days. B) Investigation ratio calculated as time investigating the 2 objects during the test phase / (time investigating the 2 objects during the test phase + during the last 5 min). Control group (C, n=15); Prenatally‐stressed group (PS, n=15); ^*^p<0.05.

### Contextual discrimination

The ability to differentiate two similar environments or contexts based on previously acquired memory for one of the contexts using contextual discrimination fear conditioning tests has been extensively used as a proxy for contextual pattern separation (McHugh et al., 2007; Sahay et al., 2011a; Kheirbek et al., 2012b; Nakashiba et al., 2012; Tronel et al., 2012). Overall, in our paradigm adapted from previous studies (Lee et al., 2011; Kheirbek et al., 2012b; Tronel et al., 2012), mice were exposed to contextual fear conditioning in a specific context (A) and we tested their ability to discriminate this context from a very similar one that shared multiples features (context B) or from a dissimilar one (context C) (Figure 2A). In context A, the test room was lit with a neon light (600 lux) and the conditioning chamber (30 × 24 × 22 cm) was lit with an overhead white light; its walls were made of clear Plexiglas; the floor of the chamber consisted of 42 stainless steel rods, separated by 3 mm, which were wired to a shock generator and scrambler (Imetronic, Pessac, France). The chamber was thoroughly cleaned with a 1% acetic acid solution between each mouse. For context B (similar context), the test room was still lit with the neon light (600 lux), the chamber (30 × 24 × 22 cm) was not lit, and the two sidewalls were covered with black panels cued with black and white patterns; the front and rear walls as well as the floor remained the same as in context A and a 1% acetic acid solution was also used to clean the chamber between each mouse. For context C (dissimilar context), the test room was lit with a fluorescent light (40 lux), the chamber (30 × 24 × 22 cm) was not lit, the side walls were made of dark grey Plexiglas, and the grid floor was covered with a plastic laminated sheet. A 5% NaOH solution was used to clean the chamber between each mouse.

**Figure 2:**
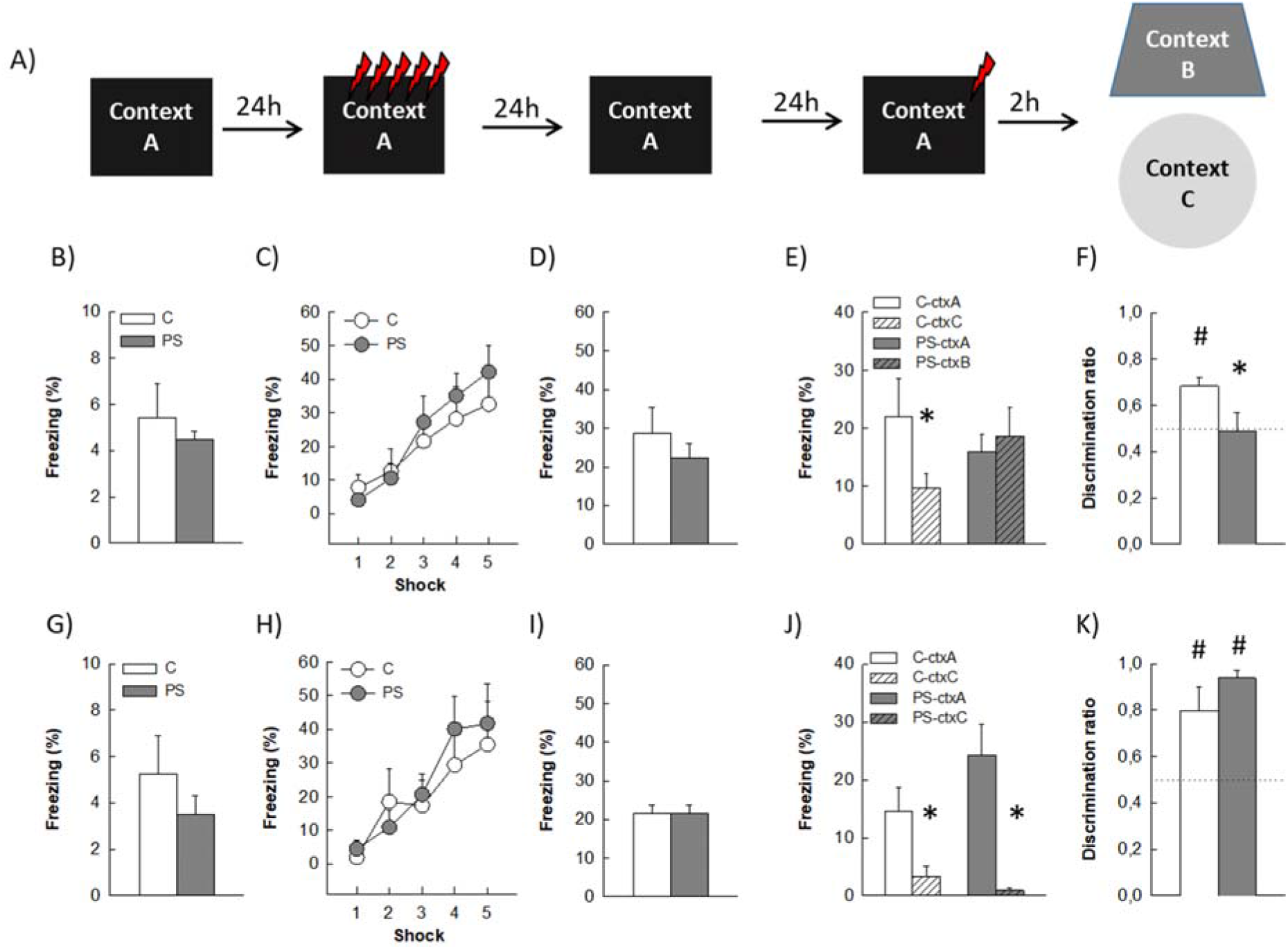
Prenatal stress alters mice abilities to discriminate similar contexts. **A)** Experimental design of the task. **B) to F)** freezing responses of mice exposed to the similar contexts A and B; **G) to K)** freezing responses of mice exposed to the different contexts A and C. **B) G)** baseline freezing levels of C and PS mice during pre‐exposure to a neutral A context. **C) H)** freezing response to the conditioning footshocks. **D) I)** freezing behavior recorded upon re‐exposure of mice to the conditioning context A. **E)** freezing levels of C and PS mice exposed to the conditioning context A (ctx A) and to a similar context B (ctx B). **J)** freezing levels of C and PS mice exposed to the conditioning context A (ctx A) and to a different context C (ctx C). **F)** Comparison of discrimination ratios calculated as freezing in ctxA / freezing in ctx A + ctx B between C and PS mice. K) Comparison of discrimination ratios calculated as freezing in ctxA / freezing in ctx A + ctx C between C and PS mice. Similar contexts groups (C n=9; PS n=13); Different contexts groups (C n=6; PS n=6); ^*^ different from control at p<0.05; # different from 0.5 at p<0.05.

On day 1 mice were pre‐exposed to context A for 10 min, and their baseline freezing behavior measured over the entire session. On day 2, they were allowed to freely explore context A again for 5 min upon which five foot shocks (1 mA, 1 s) were delivered at 90 s intervals. Freezing behavior was measured in the 90 s intervals after every consecutive foot shock. On day 3, the conditioned mice were tested in the same context for 5 min without any foot shock and their freezing behavior recorded. The next day, animals were assessed for their discrimination abilities. Mice were placed in context A and were delivered a single foot shock (1 mA, 1 s) after a 3 min exploration period. Fifteen seconds later, they were returned to their home cage for a 2 hours rest period upon which half the mice of each experimental group (C vs PS) were exposed to context B for 3 min and the other half to context C. Freezing behavior in both context A and context B/C was recorded for the first 3 min. of exposure. Freezing levels are expressed as % of the time period or as Discrimination ratios calculated as follows: Ratio = (freezing percentage in chamber A/(freezing percentage in chamber A + B or C) in which values higher than 0.5 indicate that mice are able to discriminate the 2 contexts.

### Statistical analyses

All statistical analyses were performed with Statistica 12.0 software (Statsoft). A two‐way repeated‐measures Anova with group (C v sPS) and separation load (low, medium, high) was performed on data collected during the metric separation task. A combination of Student t‐tests with group as main factor, of two‐way repeated measures Anova with (group x shock) or (group x context) as main factors were used for comparing freezing levels in C and PS mice. LSD post‐hoc analysis was performed when appropriate. All data are presented as mean ± SEM.

## RESULTS

### Metric separation is altered in PS mice

Behavioral pattern separation was first measured with a variation of a metric spatial‐processing task (Goodrich‐Hunsaker et al., 2008) in which animals are exposed to changes in the metric relationships between 2 objects. Three conditions with different levels of difficulty were tested (Figure 1A): an easy condition where objects relocation (55 cm) is very different from their initial position (20 cm) (pattern separation load is low), a medium condition, and a difficult one where the change in objects metric configuration (35 cm) is subtle compared to the initial position (20 cm) (pattern separation load is high). All mice displayed habituation to the objects during the sample phase of the trial as demonstrated by a decrease in investigation duration during the 20 min interval (data not shown). Analysis of the investigation ratio indicated that mice from both groups showed a decreased performance when task difficulty was increased from low to high pattern separation load (Figure 1B; load effect F_2,56_=5.924, p=0.004). Although this decrease was similar for both group (group x load interaction F_2,56_=0.098, p=0.9), PS mice showed an overall lower ability to detect changes in objects’ positions compared to C mice (F_1,28_=4.639, p=0.03), thereby indicating a deficit in metric separation abilities.

### Contextual discrimination is altered in PS mice

To test contextual discrimination, mice of C and PS groups were divided into two subgroups (Figure 2A): a similar context one (tested in similar contexts A and B; Figures 2B to 2F) allowing to measure behavioral pattern separation abilities and a different context one (tested in contexts A and C; Figures 2G to 2K) allowing to control for coarse discrimination abilities. On the first day, during context A pre‐exposure, both C and PS mice freely explored the fear arena without any significant difference in their baseline freezing levels, which remained very low (similar contexts group: Figure 2B, t_20_=0.730, p=0.47; different context group: Figure 2G, t_10_=0.948, p=0.36). During conditioning the following day, we observed as expected a significant increase in mice freezing behavior for every consecutive foot shock without any significant difference between C and PS animals (similar context group: Figure 2C, group effect: F_1,20_=0.15, p=0.69; training effect: F_4,80_=18.18, P<0.001; group X training interaction: F_4,80_=0.87,p=0.48; different context group: Figure 2H, group effect: F_1,10_=0.13, p=0.72; training effect: F_4,40_=9.20, p<0.001; group X training interaction: F_4,40_=0.49, p=0.74). When their contextual fear memory was tested in the conditioning context A twenty‐four hours later, both C and PS mice showed an increase freezing response compared to baseline but they did not significantly differ from each other suggesting that PS does not affect formation of fear memory related to contextual information (similar context group: Figure 2D, t_20_=0.90, p=0.37; different context group: Figure 2I, t_10_=‐0.01,p=0.99). Subsequently on the next day, mice from the similar context subgroups were tested in a new context B which was highly similar to A. As shown in Figure 2E, only C mice exhibited a significant discrimination across the two contexts with a higher freezing in A than in B, whereas PS mice were generalizing the two contexts (group effect: F_1,20_=0.05, p=0.81; context effect: F_1,20_=2.52, p=0.12; group x context interaction: F_1,20_=6.16, p<0.01; LSD post hoc test: freezing in A ≠ B in control mice (p=0.015) and not in PS mice (p=0.49)). As a consequence C and PS mice differed in their discrimination ratio (Figure 2F, t_20_=1.95, p=0.05), which was different from 0.5 (the 0.5 value reflects the absence of discrimination) in C mice only (t_8_=4.62, p=0.001 for C, t_13_=‐0.12, p=0.90 for PS). To determine whether the deficit in PS mice was specific to the discrimination of closely related contexts or extended to that of different contexts, which does not require pattern separation, discrimination between different contexts A and C was also tested. Whatever their life history, mice showed no deficit in distinguishing A from C (Figure 11 F; group effect: F_1,10_=0.94, p=0.35; context effect: F_1,10_=28.85, p<0.001; group X context interaction: F_1,10_=3.54, p=0.08). Consequently, discrimination ratios did not differ between C and PS mice (Figure 2K, t_10_=‐1.31, p=0.21), and were different from 0.5 for both groups (t_6_=2.92, p=0.03 for C, t_6_=14.12, p<0.001 for PS).

## Discussion

The present study demonstrates that prenatal stress has a profound negative impact on pattern separation abilities tested in adult male CD1 mice in two tasks requiring different modalities. Indeed, in our metric separation test that require animals to process metric spatial information in order to detect changes in the arrangements of objects (Goodrich‐Hunsaker et al., 2008; Chen et al., 2012), PS mice were impaired compared to control mice at all discrimination distances tested, whether the pattern separation load was high (task is more difficult) or low (task is easier). As expected performances of both control and PS mice decreased with increased complexity.

We confirmed this deficit in a contextual discrimination task based on fear conditioning in which the capacity of mice to recognize two similar contexts is tested. Results from this study indicate that whereas control mice are capable of dissociating the context in which they have received foot‐shocks from a very similar one, showing less freezing in the latest, PS mice showed the same level of response in the 2 contexts, indicating their incapacity to detect the subtle changes that differentiate the conditioning context from a similar one. On the opposite, both C and PS mice showed good performances at dissociating the conditioning context from a quite different one, indicating that the deficit observed in PS mice is restricted to similarity detection. This study also revealed that PS mice were not impaired in contextual fear memory, which is in accordance with previous studies performed in prenatally‐stressed rats (Markham et al., 2010; Wilson et al., 2013). There is however a strong discrepancy in studies analyzing fear conditioning after prenatal stress, as some studies also reported specific impairments in contextual fear memory in rats and mice (Griffin et al., 2003; Lee et al., 2011; Chavez et al., 2021). It is likely that species/strain differences, varied prenatal stress protocols and contextual fear memory‐testing paradigms can partly explain these discrepancies.

Behavioral pattern separation has been consistently shown to rely on adult hippocampal neurogenesis, whether it was tested through spatial discrimination abilities using a delayed‐non‐matching‐to‐place task in a eight arm radial maze or a touchscreen‐based task‐ (Clelland et al., 2009; Creer et al., 2010; Nakashiba et al., 2012) or through contextual discrimination abilities using a fear‐conditioning based task (Sahay et al., 2011a; Nakashiba et al., 2012; Tronel et al., 2012). As we and others have previously reported, prenatal stress is associated with a sharp decrease in cell proliferation over the course of life, leading to deficits in adult neurogenesis (Lemaire et al., 2000; Mandyam et al., 2008; Zuena et al., 2008; Lucassen et al., 2009; Belnoue et al., 2013; Ortega‐Martinez, 2015). It is thus very likely that alteration in adult neurogenesis represents a core mechanism of PS‐induced deficits in pattern separation. Interestingly, pre‐pubertal stress, which decreases adult neurogenesis as well, was also found to disrupt spatial discrimination tested in an object location recognition paradigm slightly different from ours (Brydges et al., 2018). Taken together with our results, this observation reinforces the hypothesis that altered hippocampal neurogenesis may be at the core of early life stress‐induced disruptions in pattern separation. As to the best of our knowledge this study is the only one with ours testing the impact of early life stress on behavioral pattern separation, this hypothesis awaits further consideration.

Although studies on pattern separation after early life adversity are sparse, prenatal stress has been shown to impair several other hippocampus‐dependent functions such as spatial learning and memory or object location memory (Lemaire et al., 2000; Ishiwata et al., 2005; Lee et al., 2011; Schulz et al., 2011), all abnormalities that could actually stem from the deficits in pattern separation that we observed. Indeed, learning and recalling spatial knowledge was shown to depend upon proper topological and metric spatial information processing, the lasted relying on pattern separation (Goodrich‐Hunsaker et al., 2008). Furthermore, the ability to disentangle incoming information by separating events based on their spatiotemporal similarities, as is performed in behavioral pattern separation, was proposed to serve as decreasing interference (Marr, 1971; McClelland et al., 1995; Rolls, 1996), and alteration in the sensitivity to memory interference could explain the deficits in spatial relational memory that are observed after prenatal stress in the water maze procedure (Lemaire et al., 2000). In this task, animals released from different starting points need to acquire different spatial representations of the environment in order to locate the hidden platform. Memory impairment could thus result from an increase in interference between these different representations that are juxtaposed.

In addition to explain some learning and memory dysfunctions, the altered pattern separation observed in prenatally‐stressed mice bears significant relevance for psychiatric illness. Indeed, *in utero* stress in humans has been suggested to be a major risk factor for developing post‐traumatic stress disorder (PTSD) in adulthood (Heim et al., 1997; Schwabe et al., 2012); one prominent feature of PTSD patients is their specific and paradoxical memory alteration in which co‐exist recurrent emotional hypermnesia for salient trauma‐related cues, and amnesia for the surrounding traumatic context (Brewin et al., 1996), ultimately impairing their ability to differentiate the correct predictors of threat from the false predictors in order to restrict their fear response only to the correct place and cues. As a result, these patients show excessive generalization of fear to “innocuous” environments that share some aspects of the previously confronted aversive event, provoking them to initiate inappropriate fear responses (Sahay et al., 2011b). This feature of PTSD is highly reminiscent of the behavioral profile exhibited by prenatally‐stressed mice and suggests a core involvement of pattern separation deficits, and by extension of adult hippocampal neurogenesis deficits. Although such hypothesis is currently not possible to test in humans due to the lack of appropriate tools, we hope that further analysis of these processes and links in our mice model may shed light on some of the pathophysiological mechanisms involved in the development of PTSD.

## Acknowledgment

We thank Mr C. Dupuy for providing excellent animal care. This work was supported by the Institut National de la Santé et de la Recherche Médicale, INSERM (to DNA) and the Centre National de la Recherche Scientifique, CNRS (to MK).

## RÉfÉrences

Belnoue L, Grosjean N, Ladeveze E, Abrous DN, Koehl M (2013) Prenatal stress inhibits hippocampal neurogenesis but spares olfactory bulb neurogenesis. PLoSOne 8:e72972.

Brewin CR, Dalgleish T, Joseph S (1996) A dual representation theory of posttraumatic stress disorder. Psychol Rev 103:670–686.

Brydges NM, Moon A, Rule L, Watkin H, Thomas KL, Hall J (2018) Sex specific effects of pre-pubertal stress on hippocampal neurogenesis and behaviour. Transl Psychiatry 8:271.

Calem M, Bromis K, McGuire P, Morgan C, Kempton MJ (2017) Meta-analysis of associations between childhood adversity and hippocampus and amygdala volume in non-clinical and general population samples. Neuroimage Clin 14:471–479.

Chavez MC, Ragusa M, Brooks K, Drake-Frazier C, Ramos I, Zajkowski M, Schulz KM (2021) Developmental stress has sex-specific effects on contextual and cued fear conditioning in adulthood. Physiol Behav 231:113314.

Chen Q, Kogan JH, Gross AK, Zhou Y, Walton NM, Shin R, Heusner CL, Miyake S, Tajinda K, Tamura K, Matsumoto M (2012) SREB2/GPR85, a schizophrenia risk factor, negatively regulates hippocampal adult neurogenesis and neurogenesis-dependent learning and memory. Eur J Neurosci 36:2597–2608.

Clelland CD, Choi M, Romberg C, Clemenson GDJ, Fragniere A, Tyers P, Jessberger S, Saksida LM, Barker RA, Gage FH, Bussey TJ (2009) A functional role for adult hippocampal neurogenesis in spatial pattern separation. Science 325:210–213.

Coe CL, Kramer M, Czeh B, Gould E, Reeves AJ, Kirschbaum C, Fuchs E (2003) Prenatal stress diminishes neurogenesis in the dentate gyrus of juvenile rhesus monkeys. Biol Psychiatry 54:1025–1034.

Creer DJ, Romberg C, Saksida LM, Van PH, Bussey TJ (2010) Running enhances spatial pattern separation in mice. ProcNatlAcadSciUSA 107:2367–2372.

Glover V (2011) Annual Research Review: Prenatal stress and the origins of psychopathology: an evolutionary perspective. JChild PsycholPsychiatry 52:356–367.

Glover V, O’Donnell KJ, O’Connor TG, Fisher J (2018) Prenatal maternal stress, fetal programming, and mechanisms underlying later psychopathology-A global perspective. Dev Psychopathol 30:843–854.

Goodrich-Hunsaker NJ, Hunsaker MR, Kesner RP (2008) The interactions and dissociations of the dorsal hippocampus subregions: how the dentate gyrus, CA3, and CA1 process spatial information. BehavNeurosci 122:16–26.

Griffin WC, Skinner HD, Salm AK, Birkle DL (2003) Mild prenatal stress in rats is associated with enhanced conditioned fear. Physiol Behav 79:209–215.

Heim C, Owens MJ, Plotsky PM, Nemeroff CB (1997) The role of early adverse life events in the etiology of depression and posttraumatic stress disorder. Focus on corticotropin-releasing factor. Ann N Y Acad Sci 821:194–207.

Ishiwata H, Shiga T, Okado N (2005) Selective serotonin reuptake inhibitor treatment of early postnatal mice reverses their prenatal stress-induced brain dysfunction. Neuroscience 133:893–901.

Kheirbek MA, Klemenhagen KC, Sahay A, Hen R (2012a) Neurogenesis and generalization: a new approach to stratify and treat anxiety disorders. NatNeurosci 15:1613–1620.

Kheirbek MA, Tannenholz L, Hen R (2012b) NR2B-dependent plasticity of adult-born granule cells is necessary for context discrimination. J Neurosci 32:8696–8702.

Lee EJ, Son GH, Chung S, Lee S, Kim J, Choi S, Kim K (2011) Impairment of fear memory consolidation in maternally stressed male mouse offspring: evidence for nongenomic glucocorticoid action on the amygdala. J Neurosci 31:7131–7140.

Lemaire V, Koehl M, Le MM, Abrous DN (2000) Prenatal stress produces learning deficits associated with an inhibition of neurogenesis in the hippocampus. ProcNatlAcadSciUSA 97:11032–11037.

Lucassen PJ, Bosch OJ, Jousma E, Kromer SA, Andrew R, Seckl JR, Neumann ID (2009) Prenatal stress reduces postnatal neurogenesis in rats selectively bred for high, but not low, anxiety: possible key role of placental 11beta-hydroxysteroid dehydrogenase type 2. EurJNeurosci 29:97–103.

Lupien SJ, McEwen BS, Gunnar MR, Heim C (2009) Effects of stress throughout the lifespan on the brain, behaviour and cognition. Nat Rev Neurosci 10:434–445.

Mandyam CD, Crawford EF, Eisch AJ, Rivier CL, Richardson HN (2008) Stress experienced in utero reduces sexual dichotomies in neurogenesis, microenvironment, and cell death in the adult rat hippocampus. DevNeurobiol 68:575–589.

Markham JA, Taylor AR, Taylor SB, Bell DB, Koenig JI (2010) Characterization of the cognitive impairments induced by prenatal exposure to stress in the rat. Front Behav Neurosci 4:173.

Marr D (1971) Simple memory: a theory for archicortex. PhilosTransRSocLond B Biol Sci 262:23–81.

McAvoy K, Besnard A, Sahay A (2015) Adult hippocampal neurogenesis and pattern separation in DG: a role for feedback inhibition in modulating sparseness to govern population-based coding. Front Syst Neurosci 9:120.

McClelland JL, McNaughton BL, O’Reilly RC (1995) Why there are complementary learning systems in the hippocampus and neocortex: insights from the successes and failures of connectionist models of learning and memory. PsycholRev 102:419–457.

McHugh TJ, Jones MW, Quinn JJ, Balthasar N, Coppari R, Elmquist JK, Lowell BB, Fanselow MS, Wilson MA, Tonegawa S (2007) Dentate gyrus NMDA receptors mediate rapid pattern separation in the hippocampal network. Science 317:94–99.

Meaney MJ, Sapolsky RM, McEwen BS (1985) The development of the glucocorticoid receptor system in the rat limbic brain. I. Ontogeny and autoregulation. Brain Res 350:159–164.

Nakashiba T, Cushman JD, Pelkey KA, Renaudineau S, Buhl DL, McHugh TJ, Rodriguez Barrera V, Chittajallu R, Iwamoto KS, McBain CJ, Fanselow MS, Tonegawa S (2012) Young dentate granule cells mediate pattern separation, whereas old granule cells facilitate pattern completion. Cell 149:188–201.

Ortega-Martinez S (2015) Influences of prenatal and postnatal stress on adult hippocampal neurogenesis: the double neurogenic niche hypothesis. BehavBrain Res 281:309–317.

Rolls ET (1996) A theory of hippocampal function in memory. Hippocampus 6:601–620.

Sahay A, Scobie KN, Hill AS, O’Carroll CM, Kheirbek MA, Burghardt NS, Fenton AA, Dranovsky A, Hen R (2011a) Increasing adult hippocampal neurogenesis is sufficient to improve pattern separation. Nature 472:466–470.

Sahay A, Wilson DA, Hen R (2011b) Pattern separation: a common function for new neurons in hippocampus and olfactory bulb. Neuron 70:582–588.

Schulz KM, Pearson JN, Neeley EW, Berger R, Leonard S, Adams CE, Stevens KE (2011) Maternal stress during pregnancy causes sex-specific alterations in offspring memory performance, social interactions, indices of anxiety, and body mass. Physiol Behav 104:340–347.

Schwabe L, Bohbot VD, Wolf OT (2012) Prenatal stress changes learning strategies in adulthood. Hippocampus 22:2136–2143.

Tronel S, Belnoue L, Grosjean N, Revest J-M, Piazza P-V, Koehl M, Abrous DN (2012) Adult-born neurons are necessary for extended contextual discrimination. Hippocampus 22:292–298.

Van den Bergh BRH, van den Heuvel MI, Lahti M, Braeken M, de Rooij SR, Entringer S, Hoyer D, Roseboom T, Räikkönen K, King S, Schwab M (2017) Prenatal developmental origins of behavior and mental health: The influence of maternal stress in pregnancy. Neurosci Biobehav Rev.

Weinstock M (2008) The long-term behavioural consequences of prenatal stress. Neurosci Biobehav Rev 32:1073–1086.

Wilson CA, Vazdarjanova A, Terry AV (2013) Exposure to variable prenatal stress in rats: effects on anxiety-related behaviors, innate and contextual fear, and fear extinction. Behav Brain Res 238:279–288.

Yun S, Reynolds RP, Masiulis I, Eisch AJ (2016) Re-evaluating the link between neuropsychiatric disorders and dysregulated adult neurogenesis. Nat Med 22:1239–1247.

Zuena AR, Mairesse J, Casolini P, Cinque C, Alema GS, Morley-Fletcher S, Chiodi V, Spagnoli LG, Gradini R, Catalani A, Nicoletti F, Maccari S (2008) Prenatal restraint stress generates two distinct behavioral and neurochemical profiles in male and female rats. PLoSOne 3:e2170.

